# A model of protein interactions for regulating plant stem cells

**DOI:** 10.1101/237933

**Authors:** Jérémy Gruel, Julia Deichmann, Benoit Landrein, Thomas Hitchcock, Henrik Jönsson

**Affiliations:** Sainsbury Laboratory, University of Cambridge, Bateman Street, Cambridge CB2 1LR, UK.; Department of Applied Mathematics and Theoretical Physics, University of Cambridge, Cambridge CB3 0DZ, UK.; Computational Biology and Biological Physics, Lund University, 223 62 Lund, Sweden.

## Abstract

The plant shoot apical meristem holds a stem cell niche from which all aerial organs originate. Using a computational approach we show that a mixture of monomers and heterodimers of the transcription factors WUSCHEL and HAIRY MERISTEM is sufficient to pattern the stem cell niche, and predict that immobile heterodimers form a regulatory ‘pocket’ surrounding the stem cells. The model achieves to reproduce an array of perturbations, including mutants and tissue size modifications. We also show its ability to reproduce the recently observed dynamical shift of the stem cell niche during the development of an axillary meristem. The work integrates recent experimental results to answer the longstanding question of how the asymmetry of expression between the stem cell marker *CLAVATA3* and its activator *WUSCHEL* is achieved, and recent findings of plasticity in the system.

## Introduction

The shoot apical meristem (SAM) is a dome shaped tissue located at the tip of the shoot. The stem cell niche it harbours is at the origin of all the aerial plant organs, making it a critical regulator of plant development [1]. It remains active over the lifespan of the plant, continuously providing new cells for developing organs while maintaining its shape and specific gene expression domains stable; a feat enabled by a tight homeostatic control.

This control is largely dependent on the *CLAVATA/ WUSCHEL* (*CLV* / *WUS*) negative feedback loop [2]. A dialogue between the apex of the SAM -the stem cell domain- and its centre is carried out by the transcription factor WUS and the CLAVATA signalling system. WUS is specifically expressed in a central domain; diffusing, it promotes stem cell fate at the tip of the SAM and represses differentiation at its periphery [3–6]. The CLV3 peptide is expressed in the stem cell domain, and is activated by WUS. It diffuses towards the inner layers of the meristem where, upon binding the CLV1 receptors, it represses the expression of WUS, thereby closing the feedback loop [7,8]. The antagonism between WUS and CLAVATA signalling is reflected in their perturbations: *wus* plants exhibit small or even arrested meristems, while *clavata* plants have massively enlarged meristems associated with increased organ counts.

The expression of WUS is promoted by cytokinin, making the plant hormone a major factor controlling the SAM homeostasis [9–12]. The expression domains of the enzymes catalysing the synthesis of the active hormone and its receptors stress the importance of the dichotomy between the external cell layers and the inner tissue of the meristem, which can explain the scaling of the SAM domains with its size [13].

The current spatial descriptions of *CLV3* regulation, and by extension, of the activation of stem cells, suggest a co-activation by WUS and either an apical [14] or epidermal [5,6,13,15,16] hypothetical signal, to generate an asymmetry between the stem cell domain and its main activator WUS. When it comes to the *CLV3* expression domain, the triple *hairy meristem (ham)* mutant displays a particularly puzzling phenotype [17]. In this mutant, *CLV3* is expressed in the centre of the meristem, overlapping with the expression domain of *WUS*; the apical or epidermal activation of *CLV3* seem difficult to conciliate with this observation.

The HAM transcription factors were recently shown to dimerize with WUS, with which they share multiple transcriptional targets. *HAM1* and *HAM2* were also shown to be expressed mostly within the inner tissue of the meristem [18], in a pattern reminiscent of the cytokinin receptors [13].

In the following, we explore the hypothesis that the HAM-WUS dimer represses the expression of *CLV3* away from the *WUS* domain, inspired by the reported central expression of *CLV3* in the meristem of *ham* plants. We first show experimentally that the expression pattern of *HAM1* and *HAM2* scales with the size of the meristem while remaining mainly expressed in the inner tissue layers; a pattern consistent with an epidermal repression of the two genes. We show that an activation of *CLV3* by WUS monomers together with a repression by HAM-WUS dimers is sufficient to pattern the stem cell domain while explaining the triple *ham* mutant, both using a two-dimensional representation of the meristem and a 3D tissue template generated from confocal microscopy. The resulting model reproduces the asymmetry between WUS and CLV3 expression domains and multiple experimentally described perturbations of the system. It allows for a plastic stem cell domain location and can provide an explanation for this recently observed developmental phenomena.

## Results

### The expression domains of *HAM1* and *HAM2* scales with meristem size

The expression domains of *HAM1* and *HAM2* were described in [18]. In both cases, the gene expression was markedly stronger in the center of the meristem than close to the epidermis. Since it is not known what regulates the HAM expression, we introduced perturbations to get an indication of the regulatory motif. To do so, we grew plants in different conditions, such that their SAM size would vary. Whatever the size of the meristematic tissue, the expression of both *HAM1* and *HAM2* is weak or null close to the epidermis, and relatively stronger in a large central domain in the rib meristem (Fig. 1). Primordia also appear to have a strong influence on the pattern; *HAM1* and *HAM2* expression are at their strongest where organs emerge.

**Fig 1:**
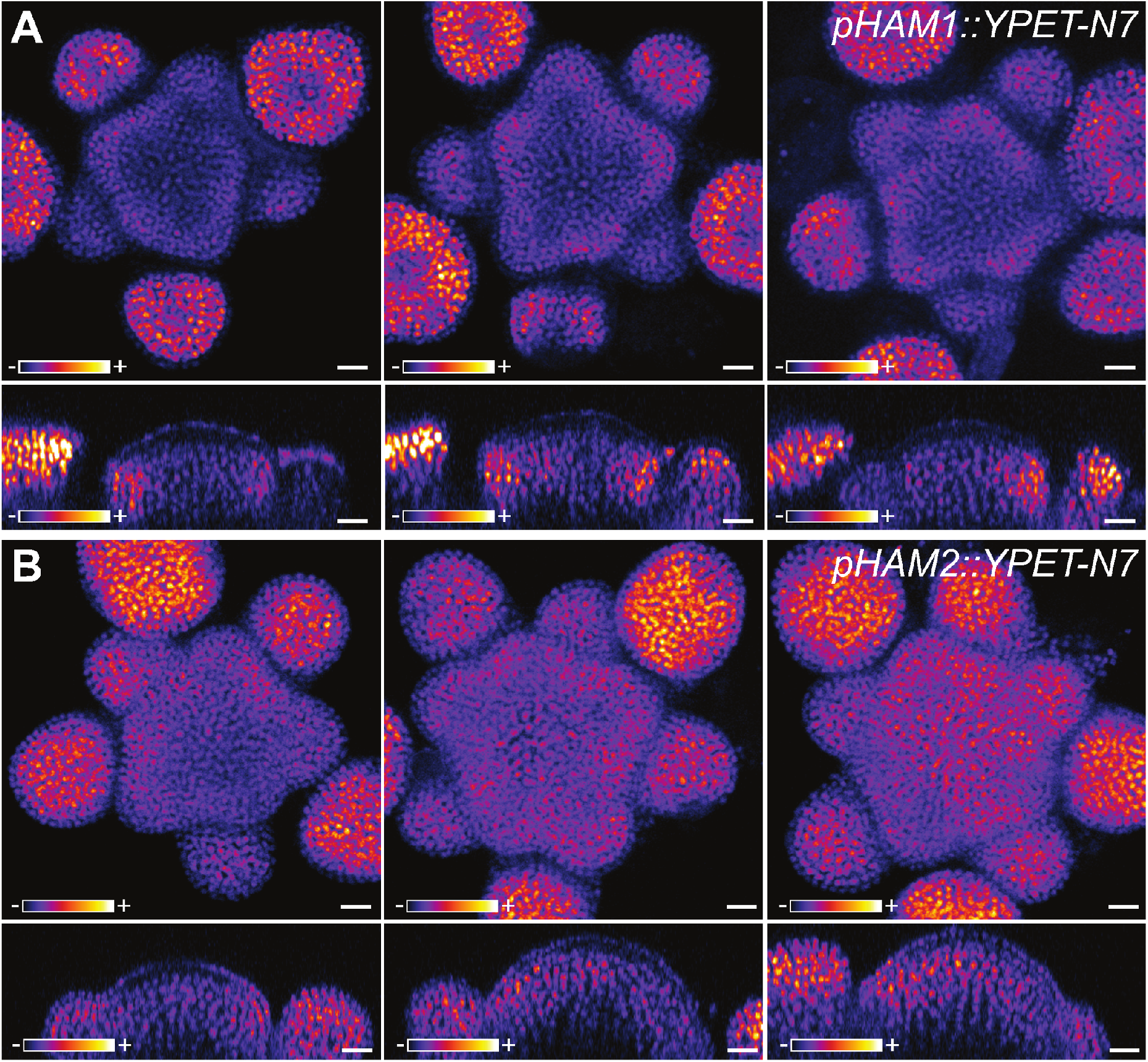
Expression of *pHAM1::YPET-N7* (A) and *pHAM2::YPET-N7* (B) in representative meristems of different sizes. Meristems were obtained by growing plants on soil, on soil supplied with fertilizer, or on a mixture of soil and sand, leading to meristems of various sizes. Top images show z-projections (sum slices) and bottom images show sections through the center of the inflorescence meristem. YPET signal is represented using the Fire lookup table of ImageJ. Scale bar: 10 μm.

The expression pattern of the two genes scales with the meristem size, while always avoiding the outer layers of the central SAM. This is consistent with a repression of *HAM* expression by a signal originating in the epidermis, as previously suggested for the cytokinin receptors [13], and this hypothesis for *HAM* regulation will be further explored in the following.

### An epidermis driven model explains the expression pattern of the stem cell domain

Given the *HAM* domains spatially overlap with WUS, a WUS-HAM dimer would be expected to have its concentration peak in the *WUS* domain and would therefore be an excellent candidate to explain the exclusion of *CLV3* from *wild type* (WT) meristems, as opposed to *ham* meristems. However, explaining the apical expression of *CLV3* with a combination of two factors expressed directly below is less straightforward, and to explore the hypothesis that HAMs together with WUS, *via* their physical interaction, are able explain the patterning of the stem cell niche in the SAM, we developed a differential equation model for the spatial expression domains.

In the model (Fig. 2A), *WUS* expression is regulated by two epidermis-originating morphogens; cytokinin acts as an activator while a second signal abstracting the repression of cyotokinin receptors from the L1 acts as an inhibitor [13]. As suggested by the scaling of *HAM1* and *HAM2* with the SAM size (Fig. 1), a third epidermis originating signal represses the expression of *HAMs* and *HAMs*, due to their functional redundancy [17], are considered as a single entity. WUS monomers can dimerize with HAM monomers, proteins undergo passive diffusion-like transport (between cells) in the tissue [5,19]. Finally, WUS monomers activate the expression of *CLV3* while WUS-HAM heterodimers repress the expression, and CLV3 peptides can move in the tissue and repress the expression of *WUS*.

**Fig 2:**
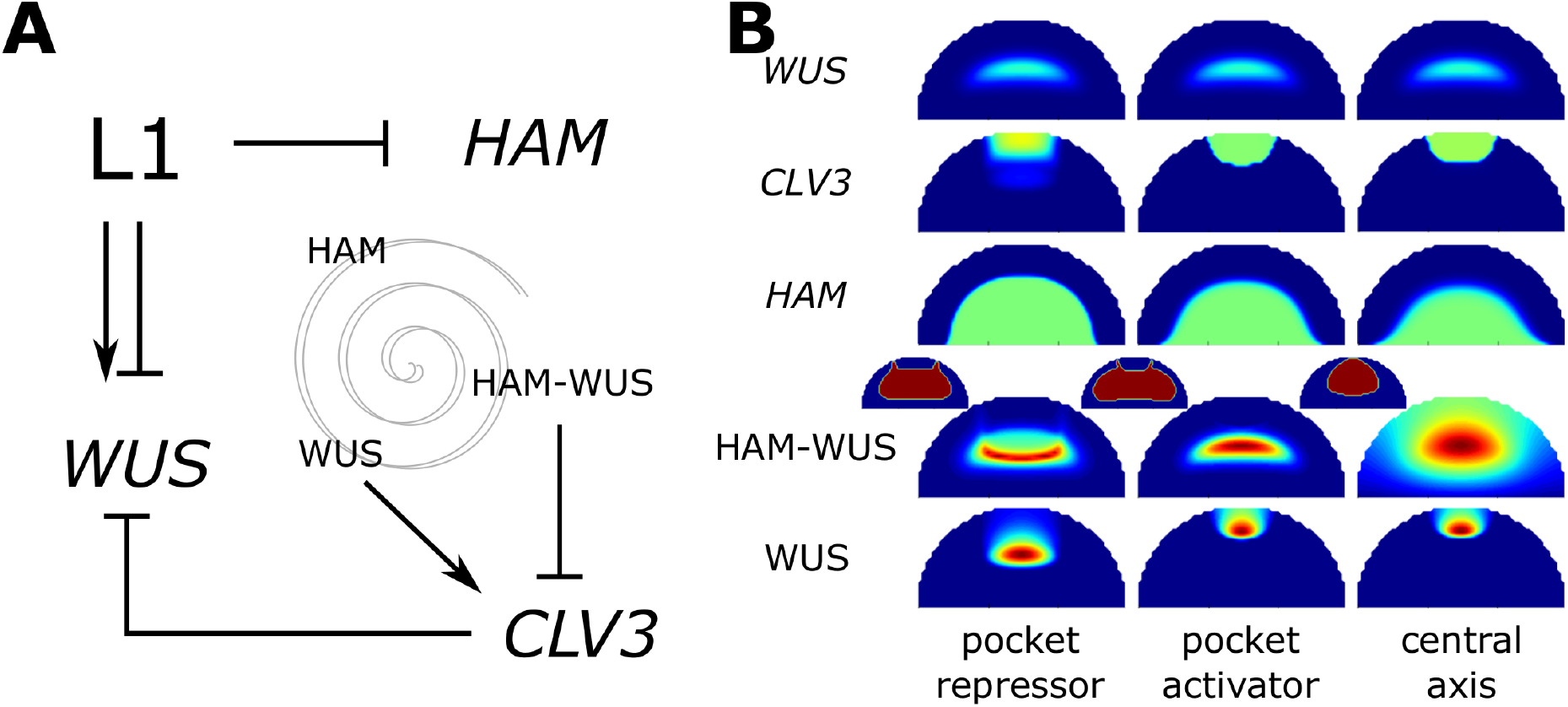
An epidermis controlled model can pattern the stem cell niche. A) Schematic of the model: the epidermis controls *WUS* expression with an incoherent feed-forward motif, it also represses the expression of *HAM.* HAM and WUS monomers can heterodimerize; WUS monomer induces the expression of *CLV3* while the heterodimer represses it. Finally the CLV3 peptide represses the expression of *WUS.* B) Parameter optimisations result in three possible behaviours. The panel presents the expression of WUS, *CLV3* and *HAM* on a color scale varying from blue (no expression) to green (optimisation target expression) to red (twice the target expression). HAM-WUS dimers and WUS monomers are plotted on a relative color scale varying from blue (minimal concentration) to red (maximal concentration). Additionally the outline of the pocket formed by the HAM-WUS dimers is displayed (smaller meristems), in blue are concentrations below the maximal value found in the epidermis and in red are concentrations above this value.

The SAM geometry is described as a half disk, the curved part of its perimeter representing the epidermis and its flat part the connection with the plant stem. This representation allows for simple deformations to mimic SAM size and shape dynamics upon mutations or growth condition variations. Model parameters are optimised to reproduce manually defined domains approximating experimental expression for WUS, *HAM* and *CLV3* at equilibrium (Materials and methods, Fig. S1).

Notably, the model achieves to reproduce the expression domains of WUS, *HAM* and *CLV3,* and multiple successful parameter values are found with the optimisation procedure (Fig. 2B). Three broadly different behaviours were identified in the resulting parameter sets. In two of them, the HAM-WUS heterodimers organize in a pocket surrounding the stem cell niche (miniature meristems in Fig. 2B). One case shows WUS monomer concentration peaking in the *WUS* expression domain, the expression domain of *CLV3* is in this case thus mostly delimitated by the repressing HAM-WUS heterodimers; this behaviour will be referred to as “pocket repressor” in the following. The second behaviour displaying a pocket of heterodimers sees WUS monomer concentration peaking in the stem cell domain, directly activating *CLV3* and the WUS activation plays a prominent role in delineating the *CLV3* domain; this behaviour is named “pocket activator”. Finally, the last scenario results in the concentration of all actors of the system peaking along the central vertical axis of the meristem and will be referred to as “central axis”. Similarly to “pocket activator”, this last case sees WUS monomers peaking in the *CLV3* domain. The three categories can be separated in planar projections of the parameter space (Fig. S2).

As all three alternative behaviours can explain the *wild type* gene expression domains in the SAM, further analysis is required to assess their biological relevance.

### The behaviour of the system hinges on protein differential mobility

First, we analysed the differences between the defined solutions in terms of predicted protein monomer and dimer mobilities. At equilibrium, the shape of the gradient formed by a diffusing molecule is controlled by the ratio between its diffusion rate and consumption rate. As described in [13], the epidermis originating morphogenes with a ratio favouring diffusion form a shallow gradient (*i.e.* cytokinin, acting at long range), while those favouring degradation fall sharply *(i.e.* the short range AHK repressor). Together these signals form the incoherent feed-forward regulation of WUS generating the adaptive scaling of the expression domain to the size of the meristem.

In the model, WUS and HAM monomers influence the mobility of each other. Indeed, if a WUS monomer is in the presence of sufficient HAM monomers, rather than diffusing, it will likely be recruited to form a dimer. This also applies to HAM monomers in the presence of a large amount of WUS monomers. As the *WUS* and *HAM* expression domains are different, the mobility of the monomers is influenced by their location within the meristem. HAM-WUS heterodimers are consumed by degradation and dissociation. The efficiency of those two reactions is not influenced by secondary species, nor is their diffusion rate; the moblility of the dimers does not vary across the tissue. We can achieve a good separation of the classes within the mobility parameter space (6 parameters), showing the relevance of this concept to compare the possible behaviours (Fig. S3).

We analysed the mobility of the two monomers and the dimer across the central axis of the SAM (Fig. 3). For each of the three categories, and as expected, WUS monomers are more mobile close to the apex of the tissue, where relatively little HAM is found, compared to close to the stem, where HAM is expressed. The opposite holds true for HAM monomers for all three categories as HAM monomers are more mobile close to the stem than they are at the apex. However, the three categories can be differentiated when monomer mobility is compared to dimer dynamics. In the “central axis” category, dimers are the most mobile of the three species. For the “pocket repressor” category, dimers are the least mobile species. For the “pocket activator” category, the dimer mobility is between the maximum and minimum mobility of both dimers.

**Fig 3:**
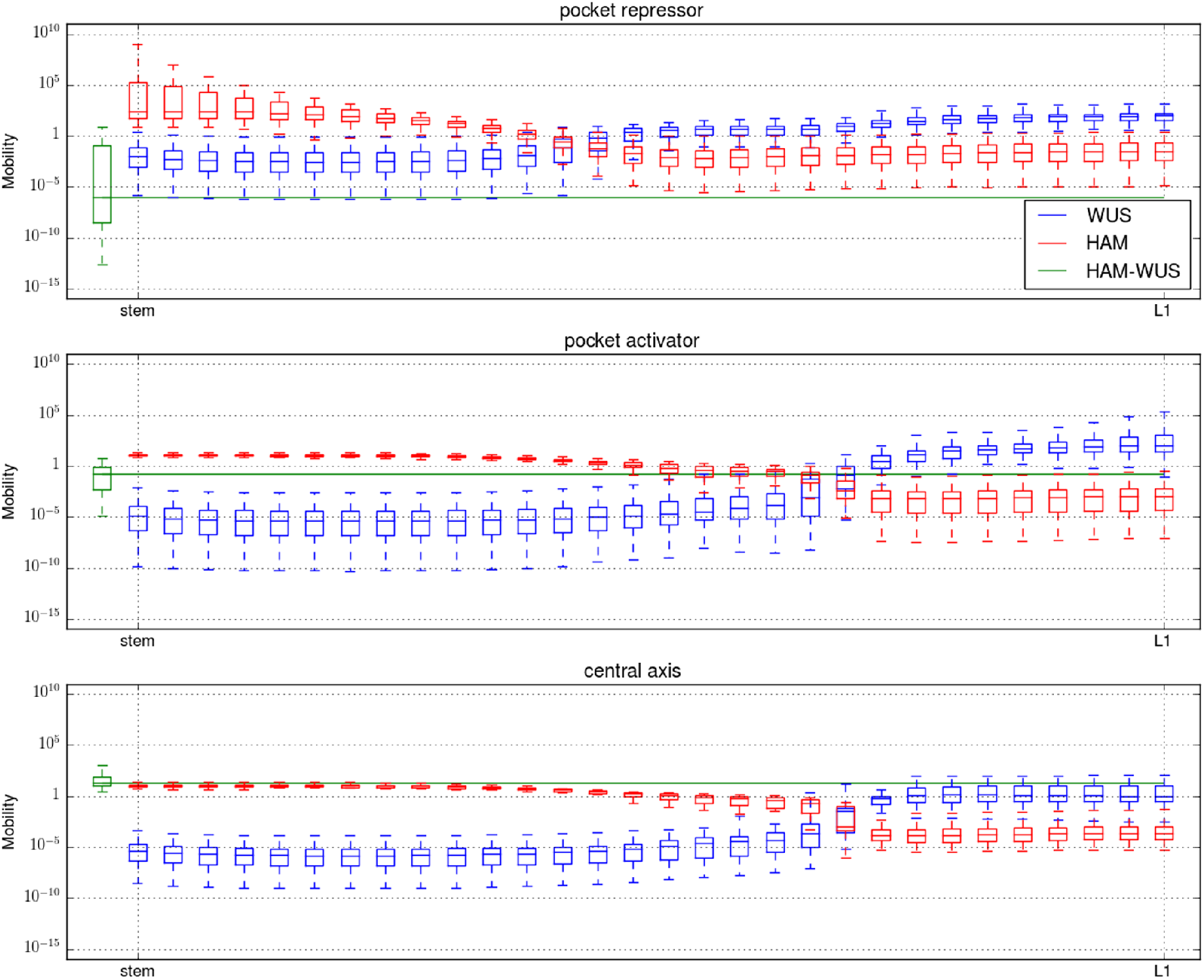
Monomer and dimer mobility. The dimer mobility is computed following 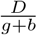 and monomer mobility following 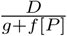 with D the diffusion rate of the considered molecule, *g* its degradation rate, *b* dimers dissociation rate, *f* dimers formation rate and [*P*] the concentration of the monomeric partner of the considered monomer. For all three panels, the *y* axis is the mobility on a logarithmic scale, the *x* axis is the central axis of the meristem, with the stem to the left and the apex to the right. For each considered molecule (WUS: red, HAM: green, HAM-WUS: blue), and each position, the mobility value is presented as a boxplot encompassing all optimised parameters belonging to one of the three parameter set categories.

It has been suggested that WUS moves between cells via the plasmodesmata, and that the size (e.g. number of GFP connected to WUS) can be restrictive for the mobility [5,19]. Given this, the fast moving dimers observed in the “central axis” category would be a less likely scenario.

### Perturbations expose behavioural differences between model categories

Next, we explored various perturbations of the system to expose possible variations in the response of the three different categories. In particular we implemented models representing *clv, ham* and *pCLV3::WUS* mutants along with changes of the tissue size. These perturbations are selected for having interesting changes to *CLV3* and *WUS* expression domains and characteristic changes in meristem size and shape (Fig. 4).

**Fig 4:**
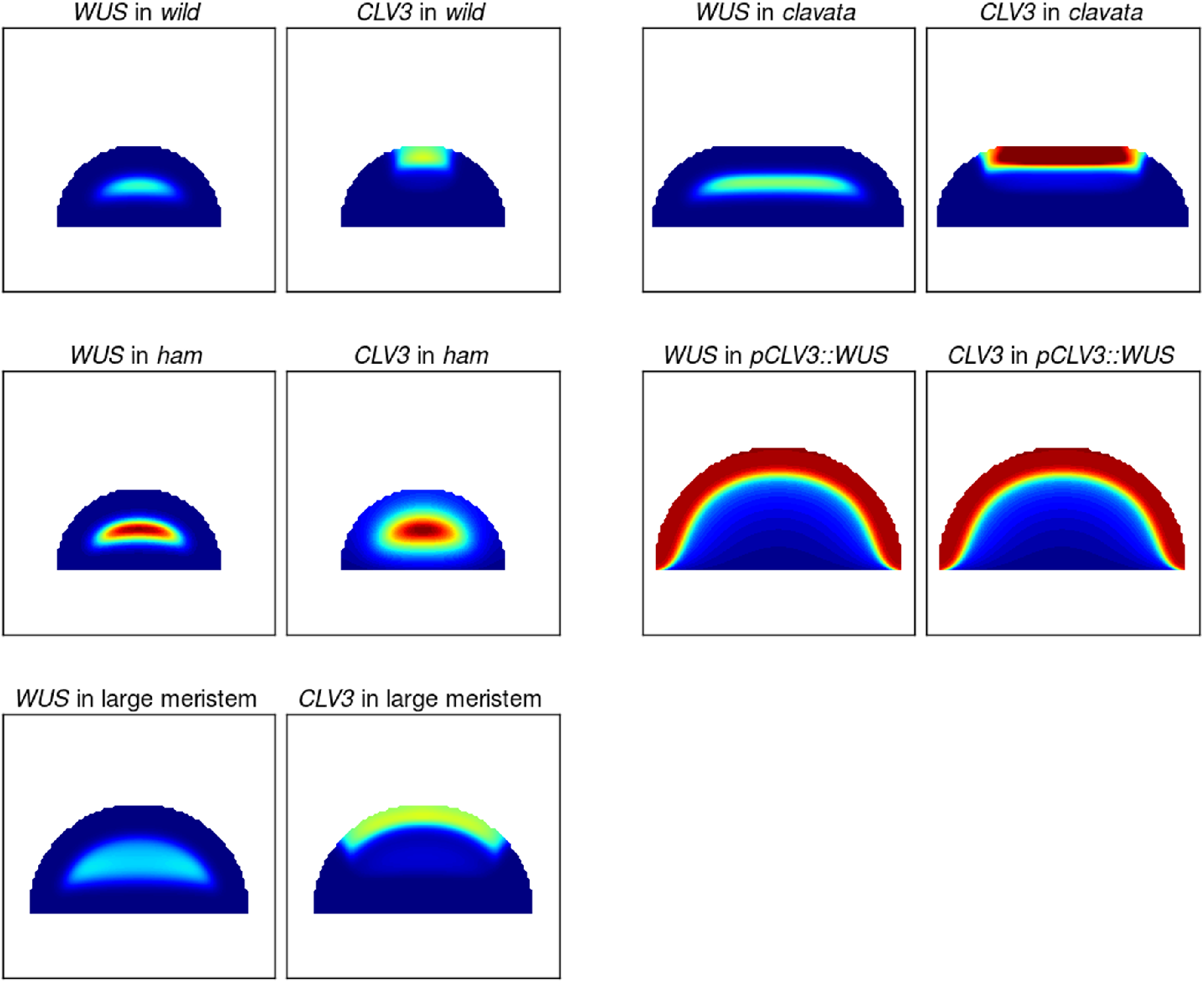
Perturbation examples. Each panel plots either the expression of *WUS* or *CLV3* in various scenarios (*wild type, clavata, ham, pCLV3::WUS* and large meristem) using an example parameter set from the “pocket repressor” category and where parameters have been adjusted to represent the mutants (Material and Methods). The colour map for *wild type, clavata* and large meristems is the same, varying from blue (no expression) to red (twice the *wild type* target expression for optimisations (green) and any value above). The colour map in the *ham* and *pCLV3::WUS* scenarios varies between blue and red (minimum and maximum gene expression in the considered settings).

A perequesite for any model aiming at describing the stem cell dynamics of the shoot apical meristem is the ability to reproduce the *clavata* mutants. Abolishing the feedback between *CLV3* and *WUS* yields fasciated meristems showing a reorganisation of gene expression domains; *CLV3* and *WUS* are expressed in two band like domains spanning the entirety of the upper cell layers of the expanded tissue [13,20] (with *CLV3* expression in the 3 to 4 first cell layers and *WUS* expression from the 3rd cell layer, the genes notably overlap in the third cell layer).

To investigate the mutant behavior across parameter values for the three categories, the expression of *CLV3* and *WUS* in WT and in a *clavata* fasciated meristem was analysed along the central axis (Fig. 5). Little differentiates the three categories, and all display an increase of the expression of both genes. If analysed in more detail, the spatial overlap of the two gene expression domains is however generally larger in the “central axis” and “pocket activator” categories (high *CLV3* expression through more than half the *WUS* domain) than it is for “pocket repressor”, where the latter category presents on average a behaviour closer to observations [13].

**Fig 5:**
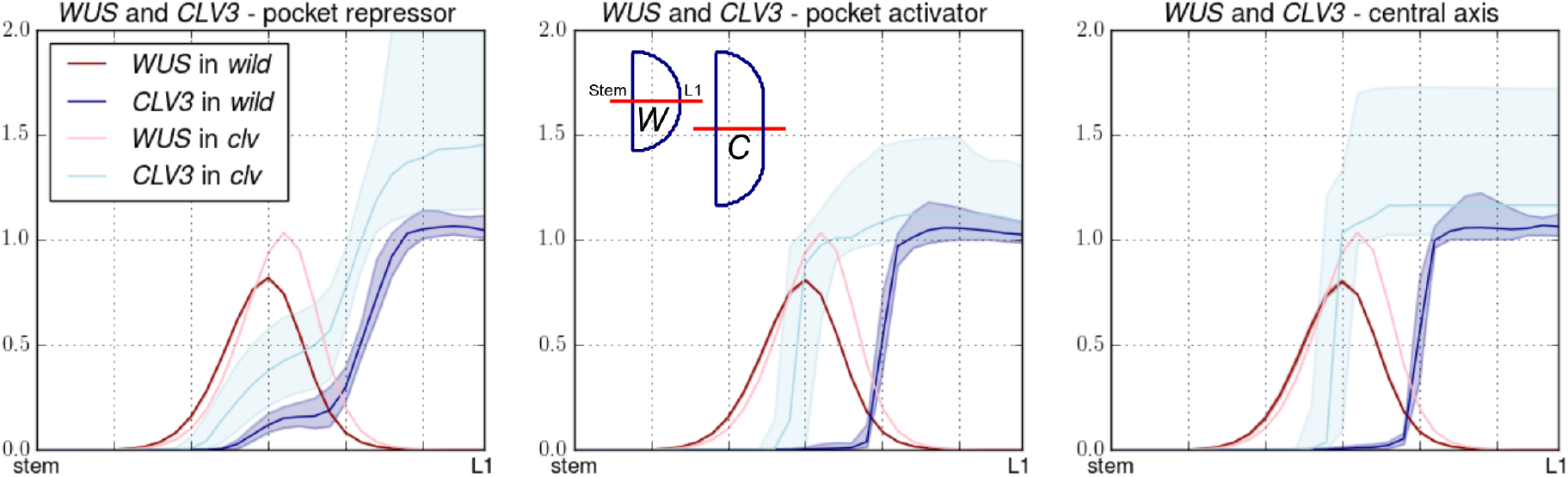
*Wild* and *clavata* meristems. The expression of *WUS* and *CLV3* along the central axis of the meristem are plotted for *wild type* and *clavata* scenarios. Each plot displays the median expression of the genes along with the first and last quartiles. For comparison, the central panel displays the shape of *wild* (W) and *clavata* (C) meristems, their central axis is marked with a red line.

The triple *ham* mutant phenotype, where *CLV3* and *WUS* expression overlap in a central domain [17], is another critical feature for the model to achieve. We analysed the relative expression of *CLV3* and *WUS* along the central axis of the meristem in simulations where the expression of *HAM* is null (Fig. 6). All three model categories have parameter values from the optimisation that can achieve a central expression of *CLV3.* Within each category, however, the domain is more or less broad and in some cases the domain even encompasses the whole tissue. The expression of WUS, activated by its epidermal incoherent feed-forward motif, is always expressed in a central domain. Due to the overlapping expression of *CLV3*, repression increases and *WUS* is consistently less expressed compared to *wild type.* Existing data does not suggest a massive drop of *WUS* expression in *ham* mutants, suggesting that either the mutation affects components of the system not modelled in this study (such as CLV3 receptors) or that the CLV3 effect on *WUS* expression is somehow buffered, as previously suggested [21].

**Fig 6:**
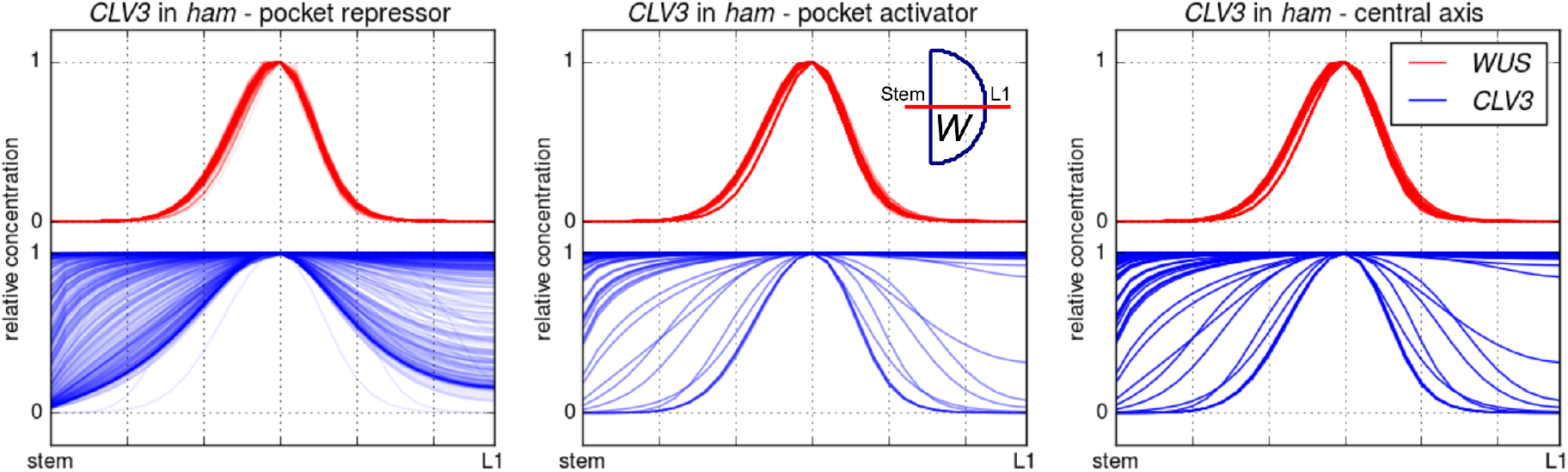
*ham* meristems. The expression of *WUS* (red) and *CLV3* (blue) along the central axis of the meristem is plotted for the *ham* scenario. Each line of each plot presents a parameter set and the gene expressions are scaled between 0 and 1 (minimum and maximum expression in the studied parameter set). Simulations were run on the *wild type* meristem shape. The transparency of the lines is adjusted in each panel for better display.

A third mutant with an interesting and non-trivial phenotype is the *pCLV3::WUS* described in [22]. Plants in which *WUS* expression is driven by *CLV3* promoter exhibit a massively enlarged meristem in which both *CLV3* and *WUS* are expressed in the three outermost cell layers of the tissue. While the direct activation of *CLV3* by an hypothetical L1 originating morphogen provides a straightforward way of achieving this behaviour [6,13], it is not obviously the case when WUS and HAM interactions control the activity of the *CLV3* promoter. The characteristic expression domains of *CLV3* and *WUS* in this mutant can also be displayed along the central axis (Fig. 7). Notably, a large proportion of the “pocket repressor” models achieve a correct representation of the experimental data, yet many of them fail the test and result in a situation where both genes are expressed at high levels across the whole tissue. The two other categories display such flat expression behaviour in a majority of the parameter values, and when they achieve an expression constrained to the outer cell layers, they do encompass more layers compared to what is seen in experimental data. Once again, the “pocket repressor” models fit data more closely than the other categories.

**Fig 7:**
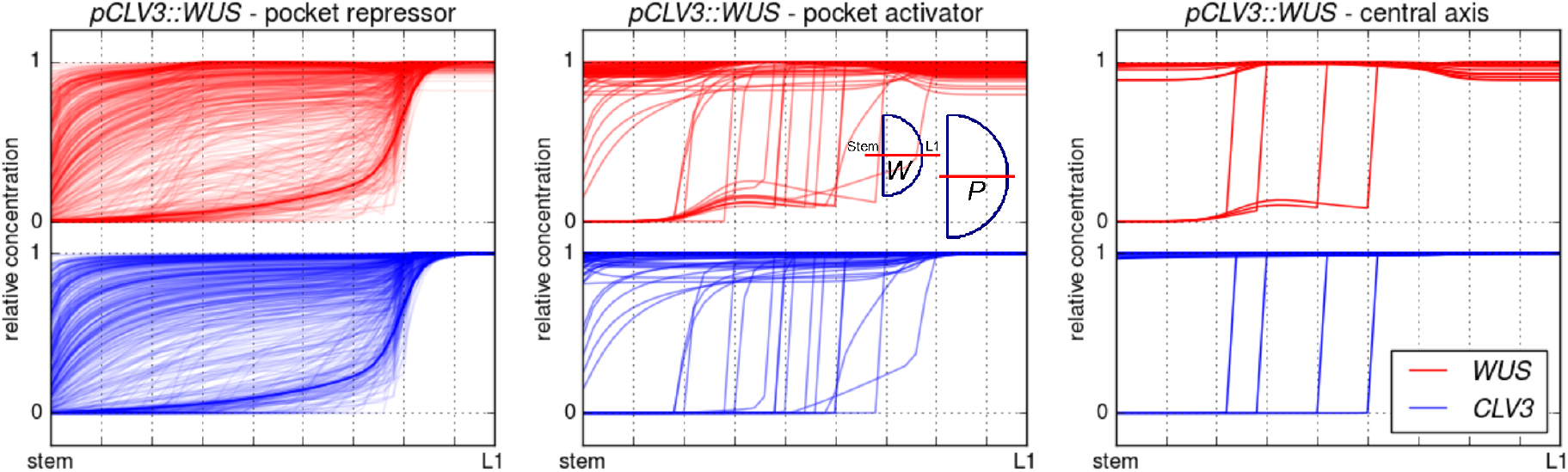
*pCLV3::WUS* meristems. The expression of *WUS* (red) and *CLV3* (blue) along the central axis of the meristems is plotted for the *pCLV3::WUS* scenario. Each line of each plot presents a parameter set and the gene expressions are scaled between 0 and 1 (minimum and maximum expression in the studied parameter set). The transparency of the lines is adjusted in each panel for better display. Simulations were run on the large meristems (P); the geometry is compared to the *wild type* (W) meristems in the central panel.

Finally, we tested the resilience of the *wild type* models to an increase of the tissue size (Fig. 8). “Pocket repressor” models behave more closely to what is seen in experiments in this situation [13], where a large majority of them maintain correct expression patterns (central *WUS* domain surmounted by an apical *CLV3* domain). In the two other categories the spatial segregation of the expression domains often breaks down, and results in a centrally expressed *CLV3* domain.

**Fig 8:**
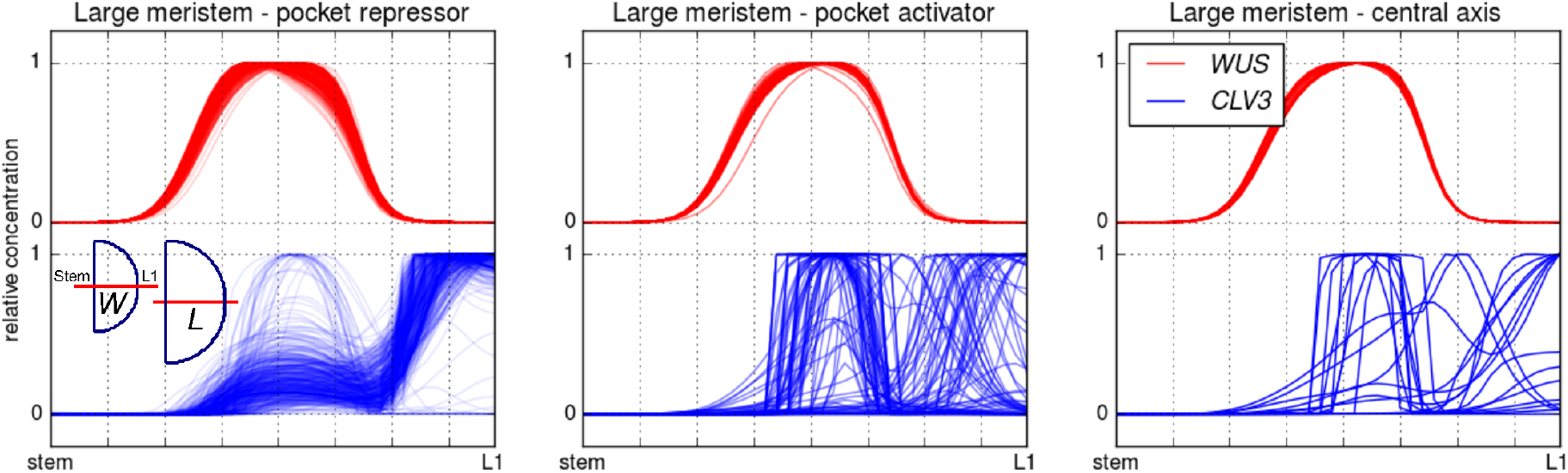
Large meristems. The expression of *WUS* and *CLV3* along the central axis of the meristems is plotted large meristems. Each line of each plot presents a parameter set and the gene expressions are scaled between 0 and 1 (minimum and maximum expression in the studied parameter set). The transparency of the lines is adjusted in each panel for better display. A comparison between large (L) and *wild type* (W) meristems is presented in the left panel.

The various mutants explored show a variety a behaviours for each of the categories, with the “pocket repressor” category generally achieving the best results. In order to assess the ability of a single subset of parameters to correctly describe all tested behaviours, we singled out the parameter sets best describing the *pCLV3::WUS* mutant (Fig. 9, designated as “pocket repressor +”). When the “pocket repressor +” parameter sets were tested against the other mutants, they also proved more successful than the rest of the category (“pocket repressor -”). We further explored the mobility of the monomers and dimers for these two categories (Fig. 10). The “pocket repressor +” parameter sets are the parameter sets exhibiting the traits separating the three original categories to the strongest extent; the dimers are the least mobile and display a large mobility differential with the monomers.

**Fig 9:**
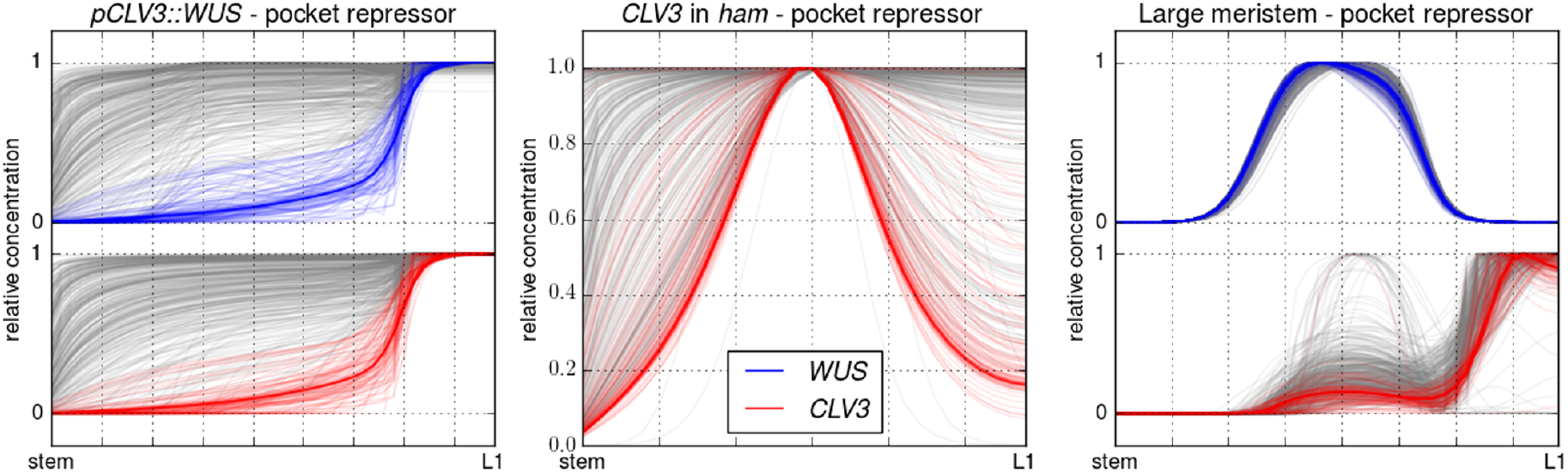
Pocket repressor perturbations exploration. The parameter sets achieving the best description of the *pCLV3::WUS* mutant are selected and coloured (pocket repressor +), leaving the rest in grey (pocket repressor -). The two sub-categories are mapped on the other two perturbations, following the same colour scheme.

**Fig 10:**
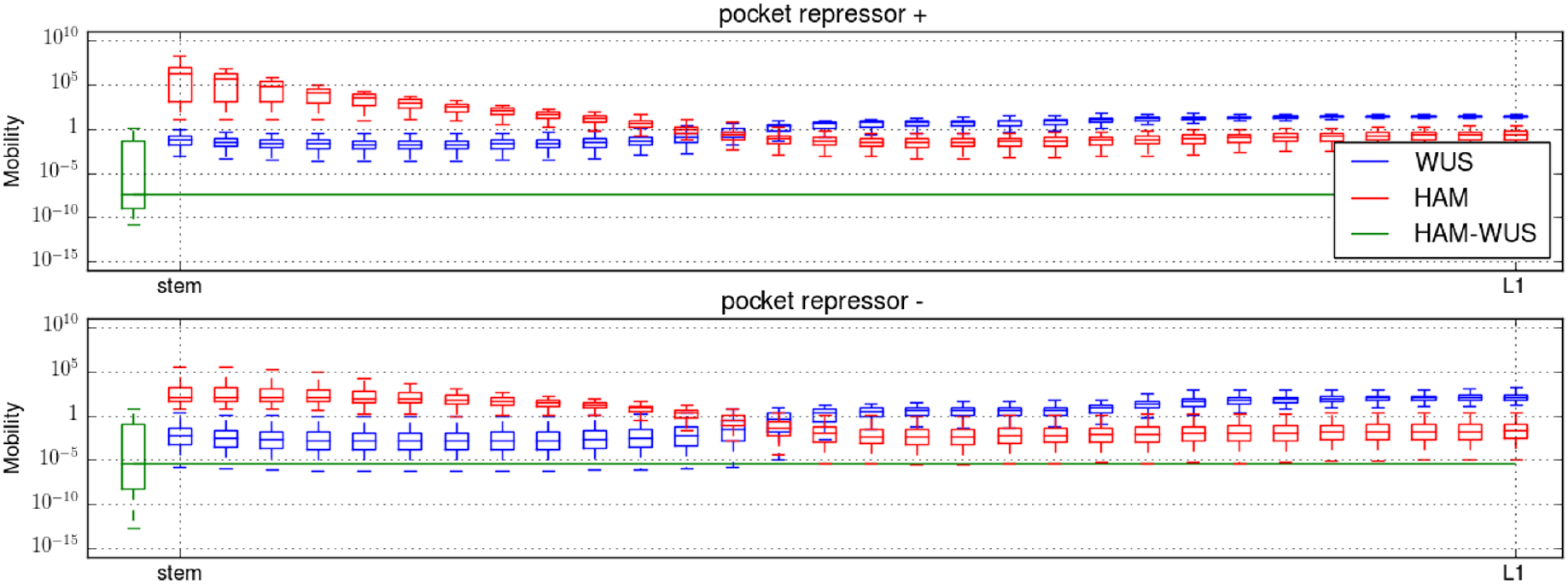
Monomer and dimer mobility in pocket repressor + and -. The dimer mobility is computed following 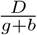 and monomer mobility following 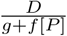 with *D* the diffusion rate of the considered molecule, *g* its degradation rate, *b* dimers dissociation rate, *f* dimers formation rate and [*P*] the concentration of the monomeric partner of the considered monomer. For all three panels, the *y* axis is the mobility on a logarithmic scale, the *x* axis is the central axis of the meristem, with the stem to the left and the apex to the right. For each considered molecule (WUS: red, HAM: green, HAM-WUS: blue), and each position, the mobility value is presented as a boxplot encompassing all optimised parameters belonging to one of the three parameter set categories.

### The position of *CLV3* shifts as HAM concentration varies

Recent studies have stressed the dynamics of *CLV3* expression, providing interesting test cases for the HAM based model. Notably, during the development of axillary meristems in the leaf axils, a *WUS* domain is established centrally in the tissue followed by an overlapping *CLV3* domain. As the meristem matures, the *CLV3* domain gradually shifts from its central position to the tip of the organ [23]. The *WUS-HAM* model can reproduce this shifting behaviour of *CLV3* expression by manipulating HAM production (Fig. 11). This suggests that during the development of axillary meristem the *WUS* and *CLV3* domains are first established, followed by a gradual expression of HAM proteins. As the *HAM* s reach their normal expression, the *CLV3* domain shifts from the center of the organ to its tip.

**Fig 11:**
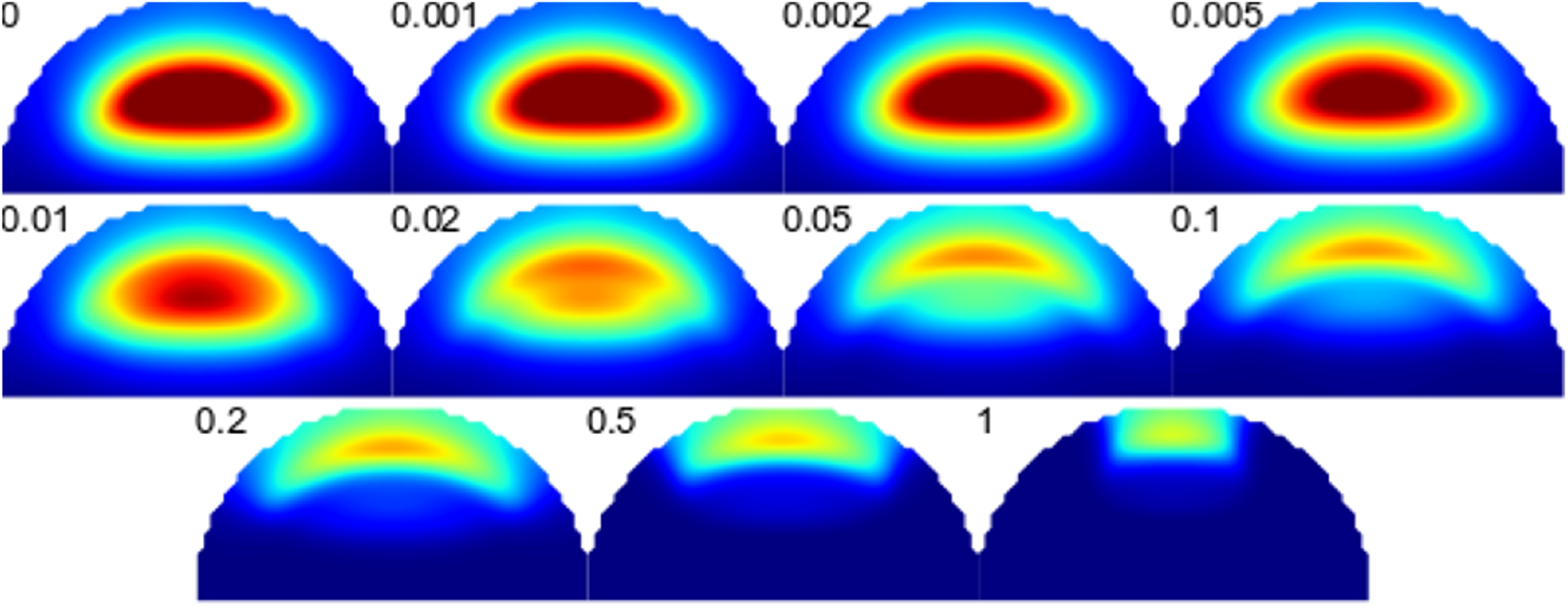
*CLV3* domain shift. For the example parameter set, the production of the *HAM* genes varies from 0 to 1, the factor is indicated along each panel. The color scale for *CLV3* expression varies from blue (null) to dark red (twice the *wild type* expression and above).

This observation also relates to recent observations made in plants with an altered *CLV3* promoter. As WUS binding sites are deleted, the expression of *CLV3* can be shifted to different locations between the apex and the the center of the meristem [24]. The authors discuss multiple hypothesis to explain the phenomenon, involving either WUS and WUS homodimers and/or possibly the involvement of the HAM proteins. Using an optimisation approach similar to the one described for the WUS-HAM model (Materials and methods), we were not able to achieve a correct patterning of *CLV3* in models exclusively using WUS monomers and homodimers when the number of dimensions of the geometrical template is higher than one (Fig. S4). This leads us to favour the HAM-WUS hypothesis, compared to a WUS-WUS hypothesis as the main regulator of *CLV3* expression patterning.

### The model is able to describe gene expression patterns on a realistic 3D meristem geometry

To confirm that a 3D geometry or meristem-specific cell neighbourhood topology does not affect the ability of the model to explain the SAM patterning, we applied the developed optimisation strategy to a 3D cell-segmented meristematic tissue [13] (Materials and methods). The description comprises cell volumes and cell contact surfaces for a meristem and early flower primordia. Optimisations resulted in models that successfully describe the gene expression of WUS, *HAM* and *CLV3* (Fig. 12), confirming that the regulatory network in the model is sufficient to generate the SAM patterns. The size of the tissue makes optimisations more difficult and prevents collecting enough parameter sets for a rigorous exploration of the parameter value space. Still, all seven obtained parameter sets belonged to category “pocket repressor”. While this analysis does not refute that it is possible to find parameter sets of the other categories, this is an indication that the “pocket repressor” category might be the best descriptor of existing data.

**Fig 12:**
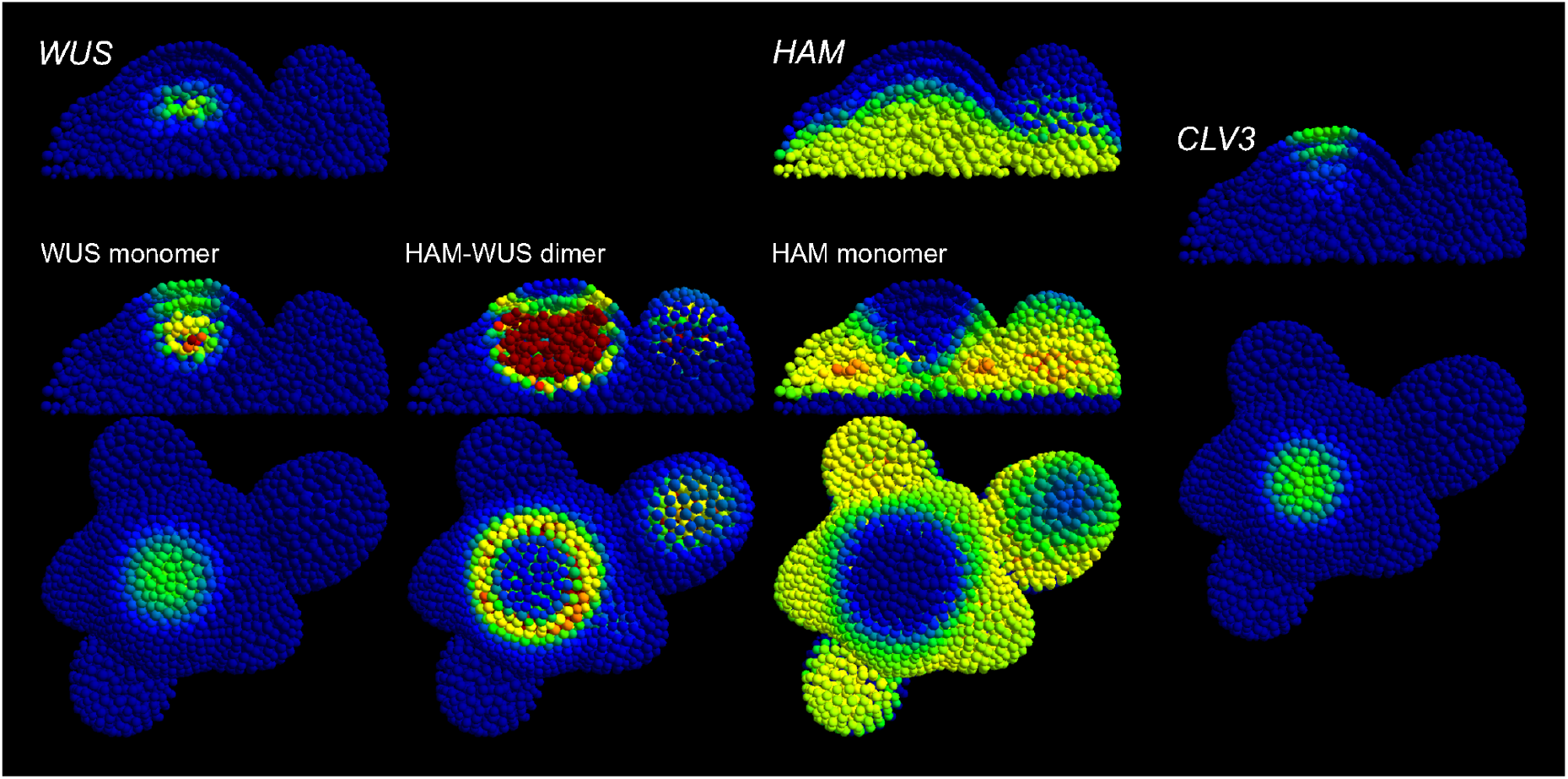
Realistic template. Example behaviour of a parameter set optimised on a realistic template. The gene expression colour map for genes is: blue for no expression, green for optimisation target expression, red for twice the target expression and any value above. The colour map for the monomers varies from blue (minimal concentration) to red (maximal concentration). In order to facilitate the visualisation of the pocket formed by the dimer, any value above the maximum dimer concentration in the L1 is displayed with red; blue indicates a low concentration.

## Discussion

In this work, we show that two genes (*WUS* and *HAM*) expressed within the inner layers of the meristematic tissue are sufficient to pattern the apical stem cell niche of *A. thaliana.* Contrary to previous studies hypothesising a direct regulation of *CLV3* by either apical or epidermal signals, the model, based on recent experimental data, achieves the plasticity required to explain the *wild type* and non-apical expression of *CLV3* observed in multiple cases: *ham* [17], mutations of *CLV3* promoter [24], lateral buds [23]. This plasticity is achieved through the double negative feedback, where the epidermis represses *HAM,* itself repressing *CLV3* together with *WUS*, and the adaptive scaling of the *HAM* expression domain itself to meristem size was confirmed in experiments where plants were grown under various nutrient conditions.

Due to the scarcity of available experimental data, we chose an exploratory strategy allowing, not only to test the potential of the model to describe the stem cell niche, but also to explore various ways of doing so. Out of three identified categories, one exhibits a more biologically probable behaviour. In the “pocket repressor” category, where the HAM-WUS dimer forms a pocket repressing the expression of *CLV3* both WUS and HAM monomers are more mobile than the dimer they form. In the “pocket repressor +” sub-category, where this diverging behaviour is even more pronounced, the model successfully represents a host of perturbations of the system, including mutants and tissue size modifications; the other categories fail to consistently achieve such results.

The main feature of the category of models able to reproduce experimental data is the repressing pocket formed by the heterodimer of WUS and HAM. While the protein mobility fits well with data on WUS, which have been shown to move between cells via plasmodesmata and where this movement is necessary for correct meristem regulation [5,19], this has yet to be confirmed for HAM proteins. Immobile HAM proteins would require a different expression pattern, closer to the repressing pocket, presently achieved *via* protein mobility.

Similarly, while WUS and HAM have been shown to have shared transcriptional targets [18], an alternative might be that HAM regulation comes solely from depleting part of the meristem from WUS monomers and hence indirectly inactivating *CLV3* expression. This can lead to a similar equilibrium regulation of *CLV3* expression, but would more closely resemble the situation of the “pocket activator” than the “pocket repressor” category of solutions, and hence direct repression by the dimer is predicted by the model. Combining the “pocket repressor” category with inactivation by WUS depletion would require the intervention of additional species to repress *CLV3* from the center of the meristem, such as WUS-WUS homodimers. This would however affect the ability of the model to reproduce *ham* mutants, where *CLV3* is expressed where the concentration of all forms of WUS would be highest.

Finally, in this model and as in [13], the main spatial readouts regulating the expression profile of cells are various signals originating in the epidermis (such as the long range diffusing cytokinin or the short range signals repressing cytokinin activity and *HAM* expression). As such, modifications of the geometry of the meristematic tissue are directly translated into modifications of the gene expression domains, possibly explaining how the meristem can adapt the size of its stem cell niche to the size of the host tissue. The handling of *HAM* in the model additionally allows it to exhibit a plastic positioning of the stem cell niche, and hence to predict recent experiments displaying such plasticity [17,23,24].

## Materials and methods

### Plant material and imaging conditions

The *pHAM1::YPET-N7* and *pHAM2::YPET-N7* reporter lines (Ler background) have been described previously [18]. Plants were grown on soil (Levington F2), on a mixture of half soil and half sand, and watered with or without 1/1/1 fertilizer (Vitafeed Standard) and placed in a constant light room (24h light, 22°C, intensity: 160μmolm^−2^ s^−1^) until bolting stage. Imaging was performed as follows: the main inflorescence meristem was cut, dissected under a binocular stereoscopic microscope to remove all the flowers down to stage 3 (as defined in [25]) and transferred to a box containing an apex culture medium (ACM) as described in [26]. Meristems were imaged in water using a 20X long-distance water objective mounted on a LSM780 confocal microscope (Zeiss, Germany). Z-stacks of 2μm spacing were taken. Z-projections (Sum slices) and orthogonal sections were performed using the ImageJ software (https://fiji.sc/).

The plant count for every conditions was:

*pHAM1::YPET-N7*:

- soil: 17 plants
- soil + fertilizer: 11 plants
- soil + sand: 20 plants

*pHAM2::YPET-N7*:

- soil: 16 plants
- soil + fertilizer: 12 plants
- soil + sand: 16 plants

### Computational methods

The following describes the differential equations defining the model, the structure used to represent the meristem as well as the methods used to find the equilibrium of the system of differential equations and to optimise model parameters. Methods for the latter are in large following previous work [13]. The section first focuses on the methodology developed to optimise the two dimensional HAM-WUS model, then we present the modifications to that methodology implemented to test the WUS-WUS model and to optimise the HAM-WUS model on the segmented meristematic tissue. Software for all the algorithms described hereafter and the resulting optimised parameter values are available http://gitlab.com/SLCU/TeamHJ/Jeremy/ham.

### 2D Meristem geometry and topology

A grid is used to represent the meristem (Fig. S1). Centered on the bottom left corner of the grid, a quarter of a circle is drawn; if the center of a grid cell is located within the circle the cell will belong to the meristem representation.

The boundaries of the meristem representation are straightforwardly extracted from this representation:

- Cells at the bottom of the grid belong to the sink, connecting the meristem to the stem of the plant.
- Cells on the left row of the grid are at located in the center of the meristem representation. This boundary is implemented as symmetric, allowing the computations to be limited to half the meristem representation.
- Cells located on the edge of the circle are the cells of the epidermis.

For a total number of cells belonging to the meristem, *n*, the sink *S* and the epidermis *L* are exported as size *n* arrays in which each cell is given a value of 1 if belonging to the said boundary and 0 if not. Similarly, each variable of the model is stored as an array of size *n*.

The neighbourhood *N* is exported as a *n* × *n* matrix where *N_ij_* = 1 if cells *i* and *j* are neighbours and *N_ij_* = 0 if not. The diagonal *N_ii_* records the amount of neighbours of i; cells belonging to the symmetric boundary have an additional neighbour representing their connection with the abstracted other half of the meristem.

### Gene expression

*WUS* and *HAM* RNA production is modelled with Hill functions. For a set of *N_A_* activators *A_a_* and *N_I_* inhibitors *I_b_*, the concentration of a RNA *X* varies as

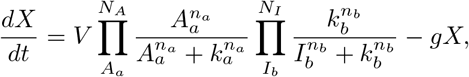

with *V*, the maximal rate of RNA production. The Hill constants *k* set the required concentration of activators or inhibitors to switch a gene between its active and inactive states. The Hill coefficients *n* control the slope of the transition between states. *gx* is the degradation rate of the RNA. The equilibrium of *X* is given by

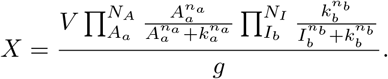

The production of *CLV3* RNA, *C*, is modelled following Shea-Ackers dynamics [27]. Considering a single binding site able to bind either WUS monomers, *w*, as activators or WUS-HAM dimers, *d*, as a inhibitors, the regulation is given by

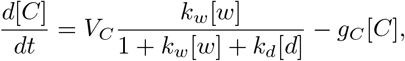

with *V_C_* the maximal rate of transcription, *k_w_* and *k_d_* the association constants for the WUS monomer and the WUS-HAM dimer. *g_C_* is the degradation rate of the RNA. The equilibrium concentration of CLV3 RNA is given by

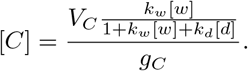

### Molecular transport

Molecular movement between cells is modelled by a passive diffusion-like transport. Such diffusing molecules are produced by a gene expression domain (*WUS, CLV3, HAM*) or the L1. This domain is referred to as *P*, a vector of cell RNA concentrations for gene expression or a vector of 1 or 0 indicating that a cell belongs to the L1 or not. The domain is associated to a production rate *p_x_* for molecule species *x*.

The bottom cell layer of the tissue represents the sink, *S*. As for the L1, it is a vector of 1 and 0. In those cells, diffusing molecules undergo degradation equal to their diffusion rate *D_x_*, approximating a continued flux into the non-modeled tissue below the meristem. Diffusing molecules also undergo a passive degradation of rate *g_x_*.

For a vector of concentration of a diffusing molecule *x*, we have

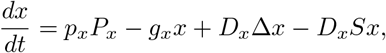

where Δ is the Laplace operator; transport in the model is assumed to be passive. For a cell *i* with *n_i_* neighbours *j*, the diffusion of *x* follows the discretised version

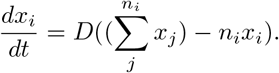

The equilibrium state of the considered molecule is found by solving

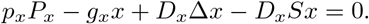

This is done with the sparse.linalg.spsolve function of the SciPy Python package.

### Model

The model describes the RNA concentration variations of *WUS (W), CLV3 (C)* and *HAM1/HAM2 (H)*, their corresponding proteins and peptides (WUS monomers (*w*), CLV3 (*c*), HAM1/HAM2 monomers (*h*), HAM-WUS dimers (*d*)) and the three positional signals produced by the epidermis (cytokinin (*L_c_*), the AHK repressor (*L_a_*), the HAM repressor (*L_h_*)). Each grid cell of the meristematic representation is described by the following system of ten equations:

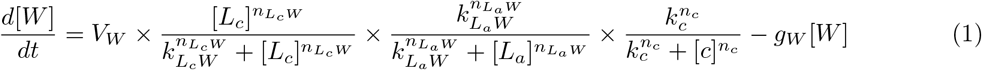

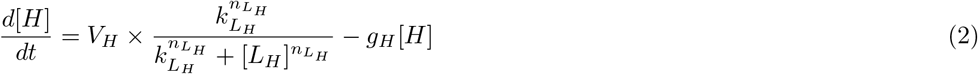

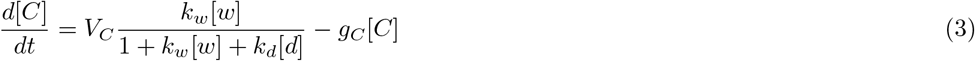

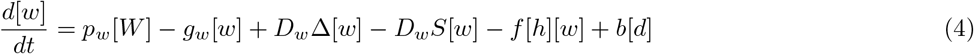

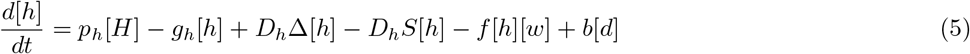

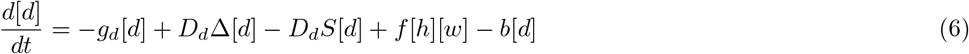

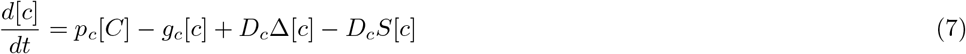

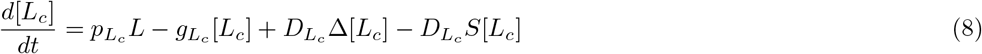

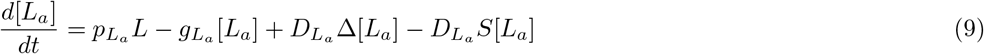

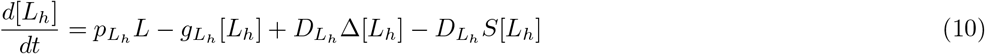

The geometric template used for optimisation includes 732 cells (i = 732), making the description of the meristem a total of 7320 equations.

### Parameter values

An multi-step model-specific optimisation strategy was designed to infer the parameter values of the model. The main difference with the strategy presented in [13] is the dimerisation of HAM and WUS that introduces additional non-linearities in the model; this step is solved with Newton’s method.

The strategy, detailed afterwards, can be summed up as follows:

1. *WUS* domain is optimised for an induced *clavata* phenotype
2. *WUS* domain is optimised for a *wild type* phenotype
3. *CLV3* and *HAM* domains are optimised for a *wild type* phenotype
4. *CLV3* and *HAM* domains are optimised for *wild type* and *CLV3* is optimised for *clavata* and *ham* phenotypes
5. CLV3 peptide is optimised to match step 2
6. the equilibrium of the whole system is computed for *wild type* and *clavata* and *ham* phenotypes

All optimisations are carried out with the L-BFGS algorithm from the SciPy Python package, the parameter values are bounded by [10^−8^ : 10^8^]. After each step described above, the subset of optimised parameters (see below) are kept as fixed values in the following steps.

In the following, diff(P) will refer to computing the equilibrium concentration of a diffusing molecule across the tissue, given a production domain P. P is a cell vector containing either ones (L1) or zeros (non-L1) if the production domain is the L1. P is a cell vector of RNA concentrations if it refers to the expression domains of WUS, *CLV3* or *HAM1/HAM2* (the equilibrium computation is described in the Molecular transport section). Similarly, *eq*([*A*_1_,…, *A_N_A__*], [*I*_1_,…, *I_N_I__*]) will refer to computing the equilibrium of a gene expression regulated by *N_A_* activators *A* and *N_I_* inhibitors *I* (both activators and inhibitors are cell vectors of diffusing molecule concentrations; the equilibrium computation is described in the Gene expression section); note that *WUS* and *HAM* are computed with Hill functions while *CLV3* uses Shea-Ackers.

Target expression domains for WUS, *CLV3* and *HAM1/HAM2* were manually defined on the optimisation template (*W_t_*, *C_t_* and *H_t_* - Fig S5). They are cell vectors containing ones (cells expressing the considered gene) or zeros (cells not expressing the gene).

#### 1) *WUS* expression domain

The first step optimises the *WUS* domain for a *clavata* phenotype; the only regulators of *WUS* here are cytokinin and the AHK repressor, both modelled as morphogens produced in the L1.

The equilibrium for the *WUS* (*W*_1_) domain is computed following:

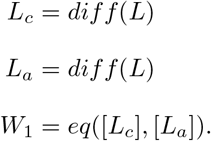

The cost function minimises the difference between the equilibrium *WUS* domain and an increased target *WUS* domain (*W_t_* × 1.5):

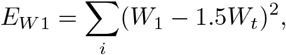

where the values for *W*_1_ and *W_t_* are from the individual cells *i*. Parameters *k_L_c__, k_L_a__, p_c_, D_c_, g_c_, p_a_, D_a_, g_a_* are optimised. As CLV3 is not considered in this step, the equation describing *WUS* dynamics is reduced to 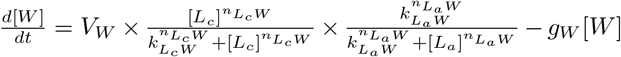. Parameters with fixed value are *V_w_* = 4, *n_L_c__* = 8, and *n_L_a__* = 4.

#### 2) *WUS* expression domain

The second step adds CLV3’s effect on *WUS*; the peptide is produced by the *CLV3* target domain. The optimisation minimises the difference between the *WUS* domain and the *WUS* target domain. The equilibrium of the *wild type WUS* domain is given by

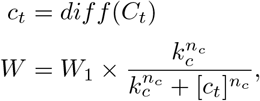

and the cost function is

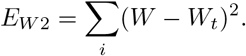

Parameters *k_c_, p_c_* and *D_c_* are optimised, and *n_c_* = 2 and *g_c_* = 1 are kept constant (together with parameters optimised in step 1).

#### 3) *CLV3* and *HAM* expression domains

This step minimises the difference between *CLV3* and *HAM1/HAM2* domains and their target domains *C_t_* and *H_t_*. After *W* and *H* equilibria are obtained, the equilibrium for the non-linear *w, h, d* sub-system can be computed using Newton’s method leading to the equilibrium for *C*.

With *F*(*x, W, H*) the 3*i* length system containing equations (4), (5) and (6) for all cells, *J*(*x, W, H*) its (3*i*)^2^ Jacobian matrix and N(*F*(*x, W, H*), *J*(*x, W, H*)) Newton’s method applied to find the root of the three-equation system, the equilibrium of *CLV3* and *HAM* is obtained as follows:

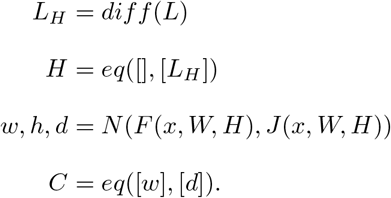

The cost function to minimise is given by

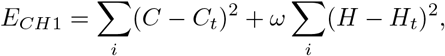

where we used a weight *ω* = 0.08. Optimised parameters are: *k_L_H__*, *p_L_H__*, *D_L_h__*, *p_w_, D_w_, p_h_, D_h_, f, D_d_, V_C_, k_w_, k_d_, g_h_, g_w_, g_d_, b*. Fixed parameters are: *V_H_* = 1, *n_L_H__* = 4.

#### 4) *CLV3* and *HAM* expression domains

This step uses the previously optimised parameter values as an initial guess to start a second broader optimisation. Here, *CLV3* and *HAM* domains are optimised for *wild type* and *CLV3* is optimised for *clavata* and *ham* phenotypes.

With *C_c_* and *C_h_* the equilibria of *CLV3* in *clavata* and *ham* phenotypes, the equilibria for the different genes and conditions are computed by the following procedure:

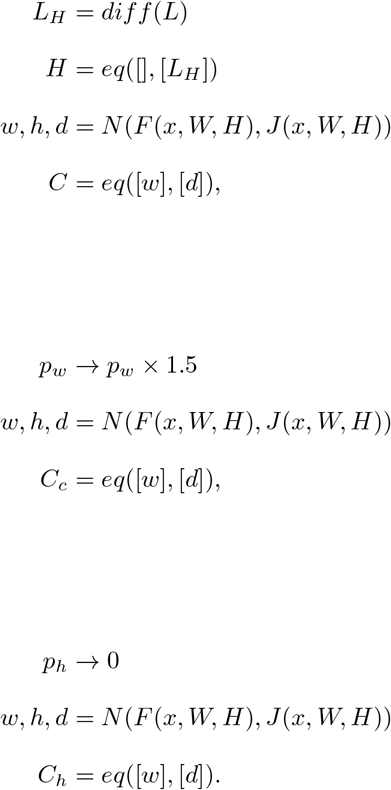

The cost function is given by:

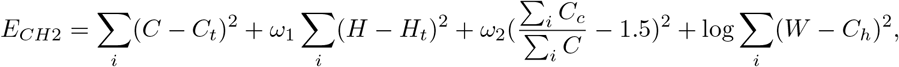

with weights *ω_i_* = 0.04 and *ω*_2_ = 0.2. Parameters *k_L_H__*, *p_l_H__*, *D_L_H__, p_w_, D_w_*, *p_h_, D_h_, f, D_d_, V_C_*, *k_w_, k_d_, g_h_, g_w_, g_d_, b* are re-optimised.

#### 5) CLV3 peptide gradient

In this step, the CLV3 peptide gradient produced by the *CLV3* domain optimised in the previous step is fitted to the gradient obtained in step 2) and produced by the target *CLV3* domain.

*C* at equilibrium is computed following:

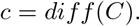

The cost function is given by

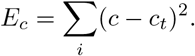

Parameters *p_c_, D_c_* and *g_c_* are optimised.

#### 6) Equilibrium of the complete model

In the final step, the previously optimised sub-parts of the model are assembled and the equilibrium of the full model is computed. The following algorithm is used: After multiple optimisation runs, the parameters kept for further analysis are those where *Σ_i_*(*C-C_t_*)^2^ < 15 when the system is in equilibrium.

### WUS-WUS model

In order to test a model in which the combination of WUS monomers and WUS-WUS homodimers would regulate the expression of *CLV3*, we used the same core model as previously described.

We removed the equations referring to *HAM* transcription and HAM monomer (equations 2 and 5). Equation 6, describing the dimer dynamics, was modified to

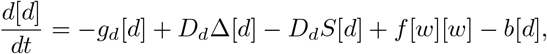

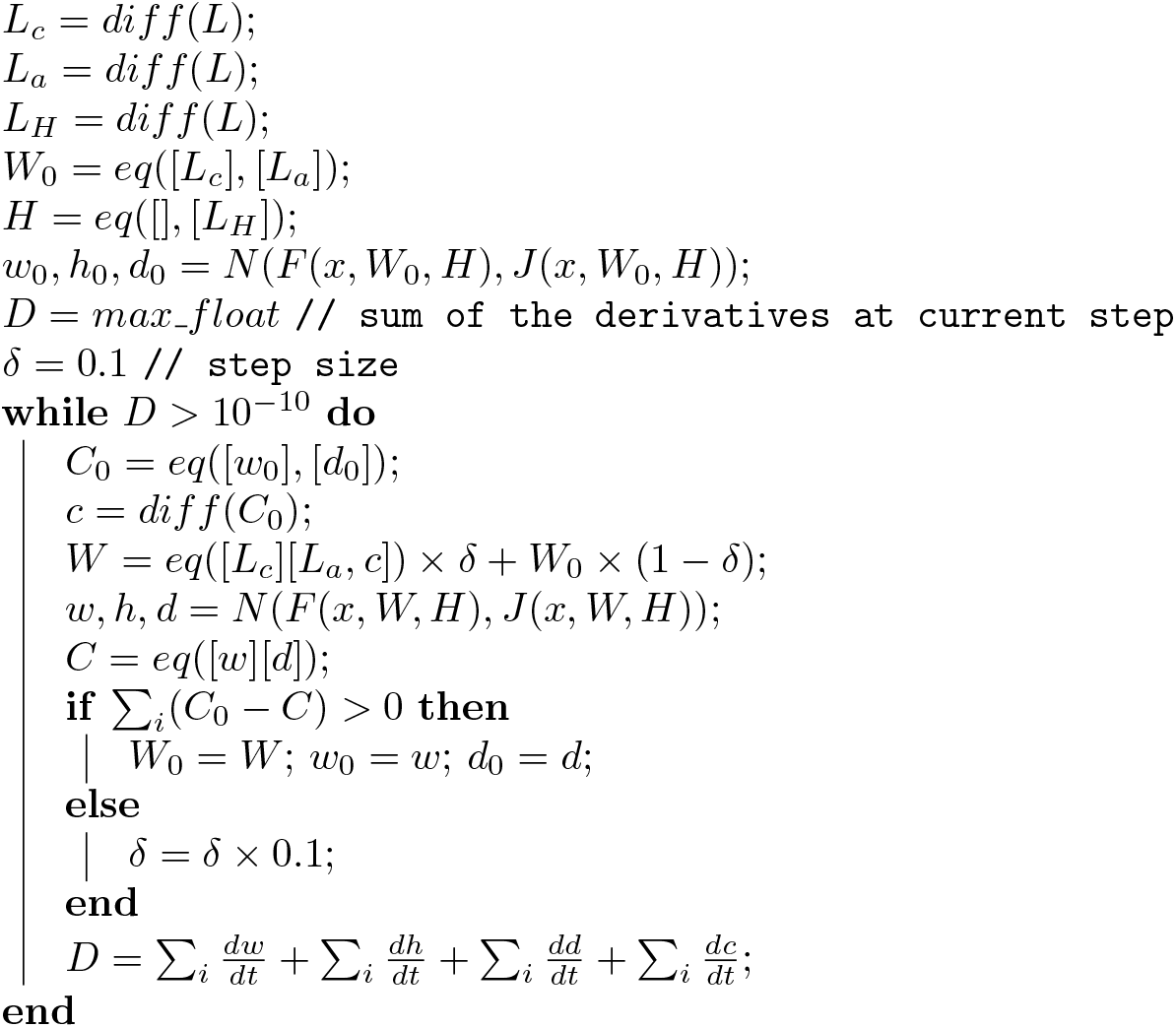

and equation 4, describing WUS monomer dynamics, was modified to

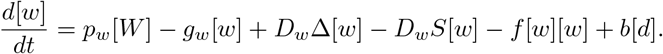

Finally the components of the optimisation procedure related to HAM expression were left out. Example results obtained with this approach are presented in Fig. S4.

### Realistic template

The main difference for the 3D template compared to the two dimensional template is the use of cell volumes and cell contact surfaces as obtained from the segmentation of the confocal imaging of a meristem tissue [13]. The equation controlling the molecular transport of x becomes:

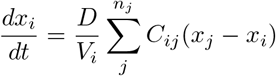

for a cell *i* with *n_i_* neighbours *j*, with *V_i_* the volume of cell *i* and *C_ij_* the contact surface between cells *i* and *j*. Hill coefficients from the optimisations in [13] were kept: *n_L_c_W_* = 7.25968619416, *n_L_α_W_* = 1.99109438845, *n_c_* = 6.66419523049, and in addition *n_lH_* = 6 was fixed. Other parameter values were found by the same optimisation strategy as used for the 2D templates.

## Acknowledgments

We thank E. Meyerowitz for providing the *HAM1/HAM2* reporter lines and members of the Jönsson and Meyerowitz (Caltech) groups for fruitful discussions. This work was supported by the Gatsby Charitable Foundation *via* grant GAT3395-PR4.

## Supporting information

**Fig S1:**
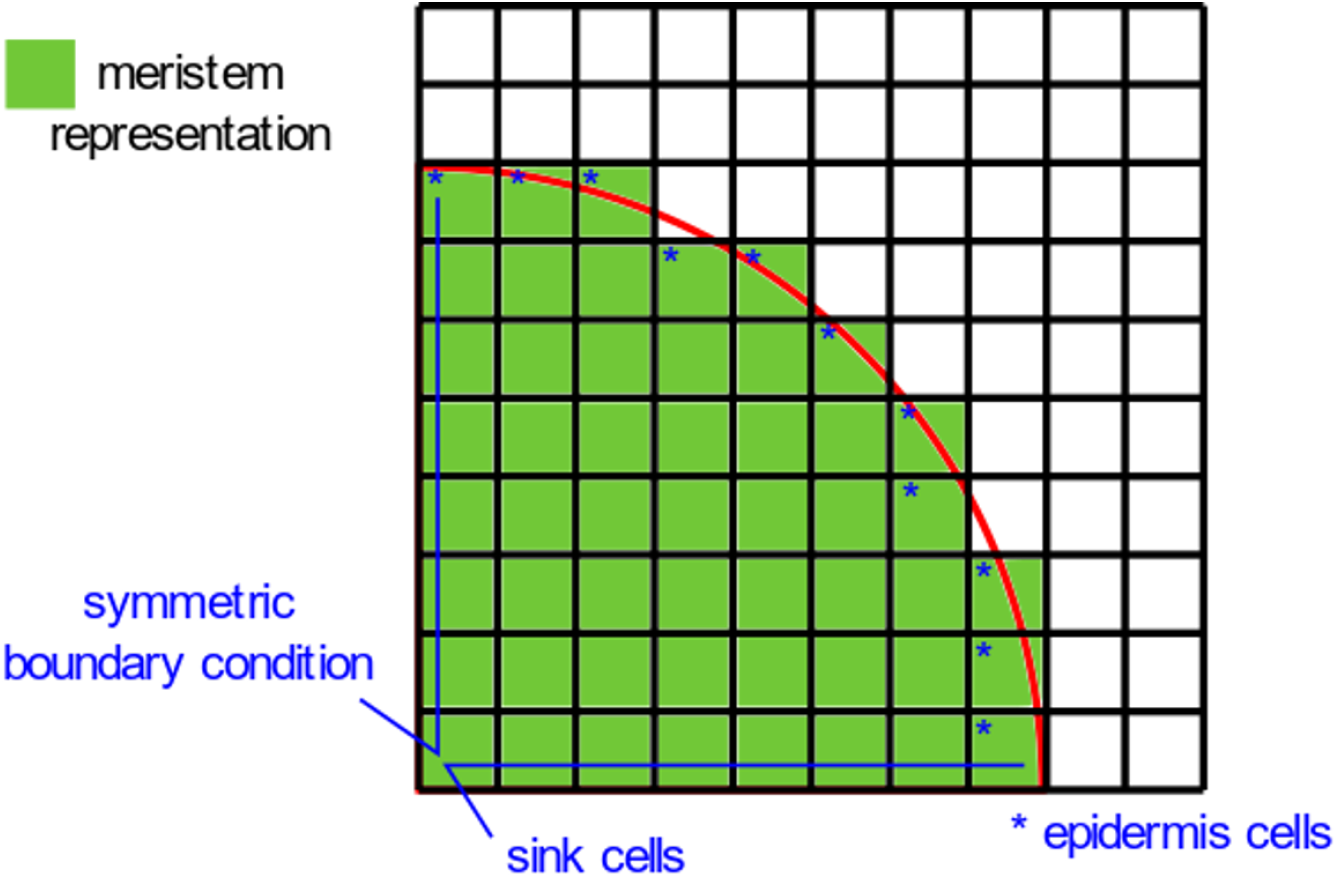
Meristem geometry layout.

**Fig S2:**
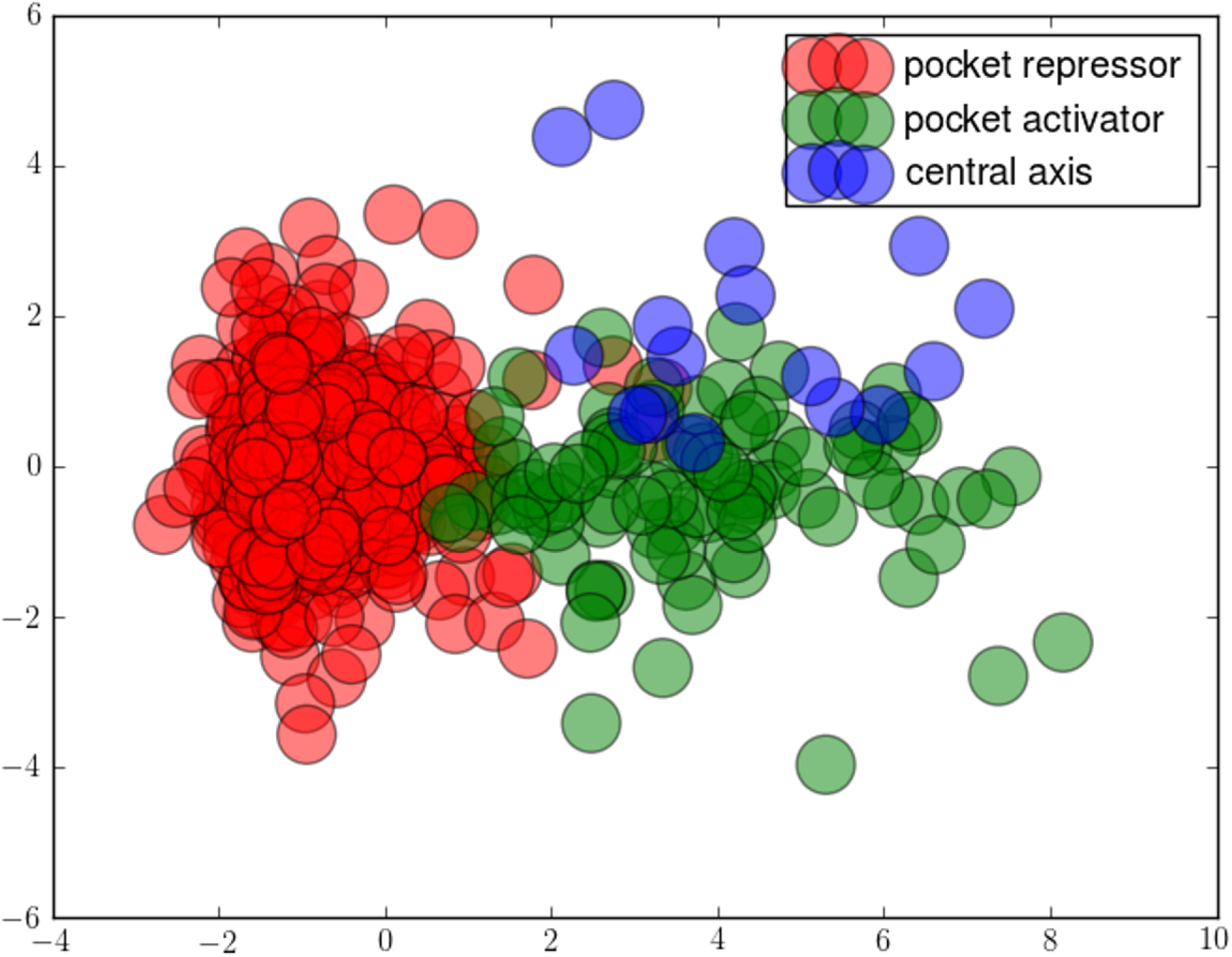
The three categories of behaviour can be separated in the parameter space. The base 10 logarithm of parameter values were centered and scaled before running a linear discriminant analysis. The figure displays the projection of the parameter sets along the most discriminative directions. The linear discriminant analysis algorithm is from the scikit-learn python package.

**Fig S3:**
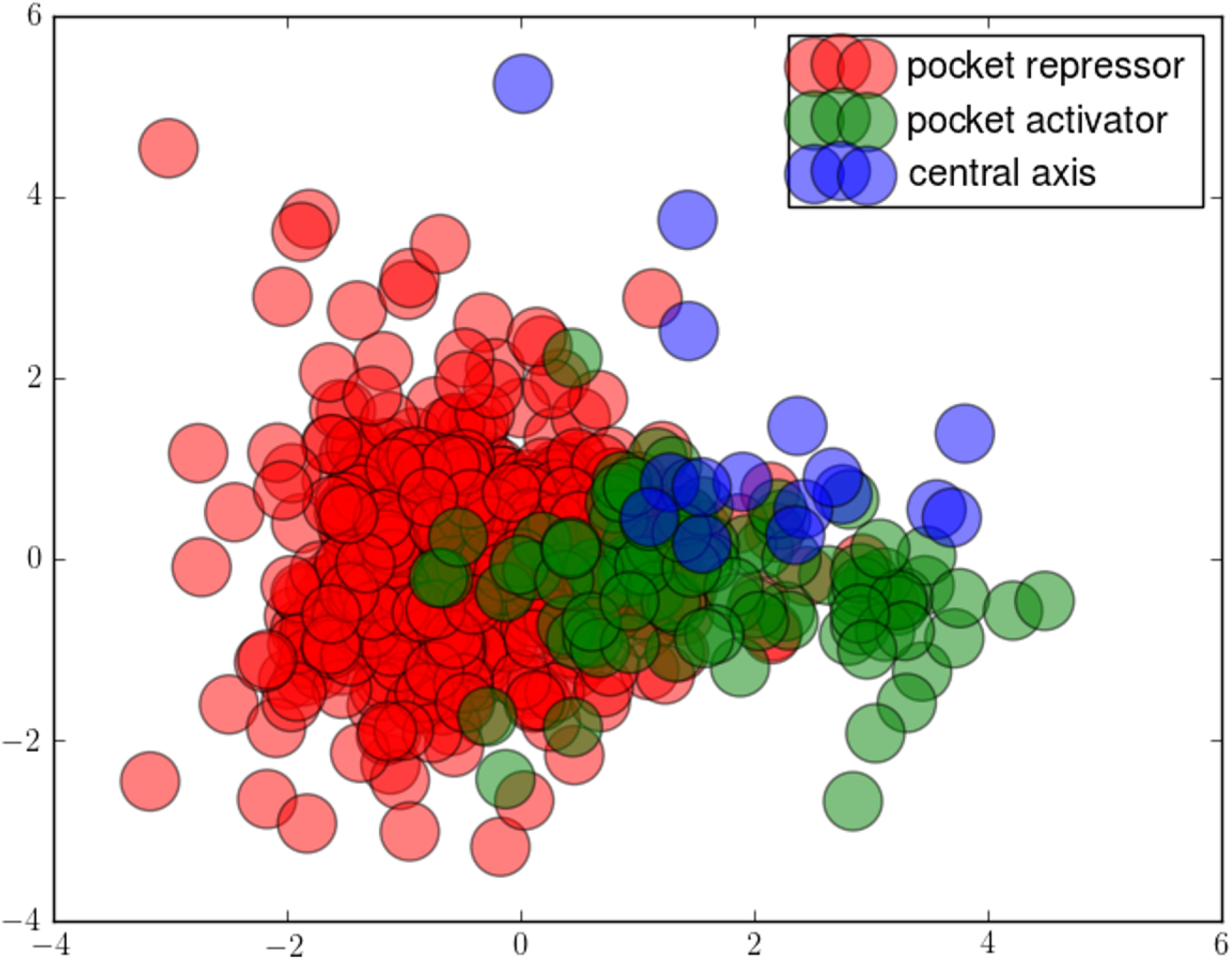
The three categories of behaviour can be separated in the parameter space of mobility parameters. The base 10 logarithm of mobility parameter values (*D_w_, D_h_, D_d_*, *g_w_, g_h_, g_d_*) were centered and scaled before running a linear discriminant analysis. The figure displays the projection of the parameter sets along the most discriminative directions. The linear discriminant analysis algorithm is from the scikit-learn python package.

**Fig S4:**
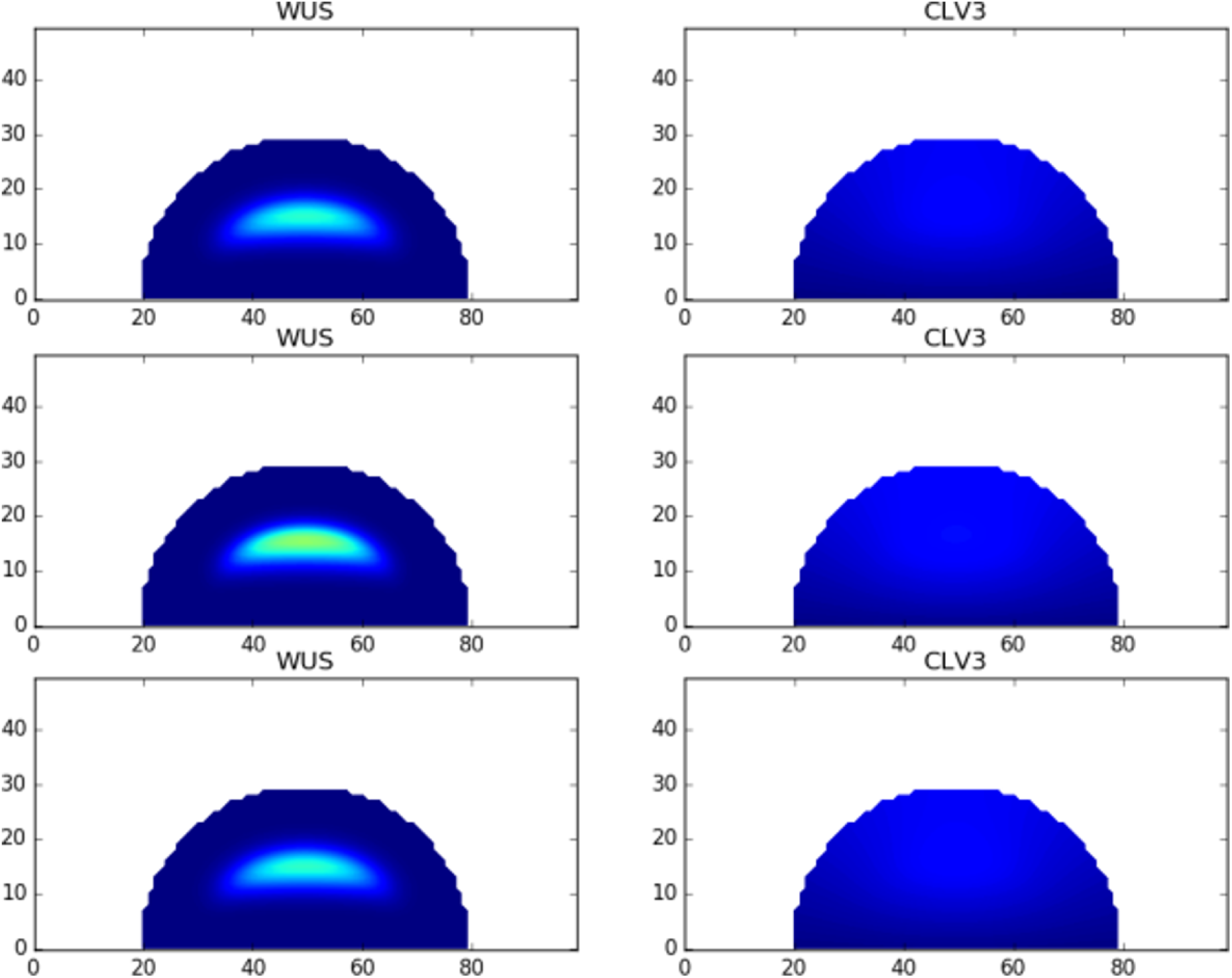
Optimisation results for a model based entirely on WUS regulating *CLV3.* The optimisation procedure was adapted to test a model where WUS is the only regulator of *CLV3* expression. The rows show three examples of the results obtained: in this scenario the best *CLV3* expression is a faint hue covering the *WUS* expression domain and the tip of the meristem. Expression of genes in the panels goes from blue (null) to red (twice the optimisation target expression and above) *via* green (optimisation target).

**Fig S5:**
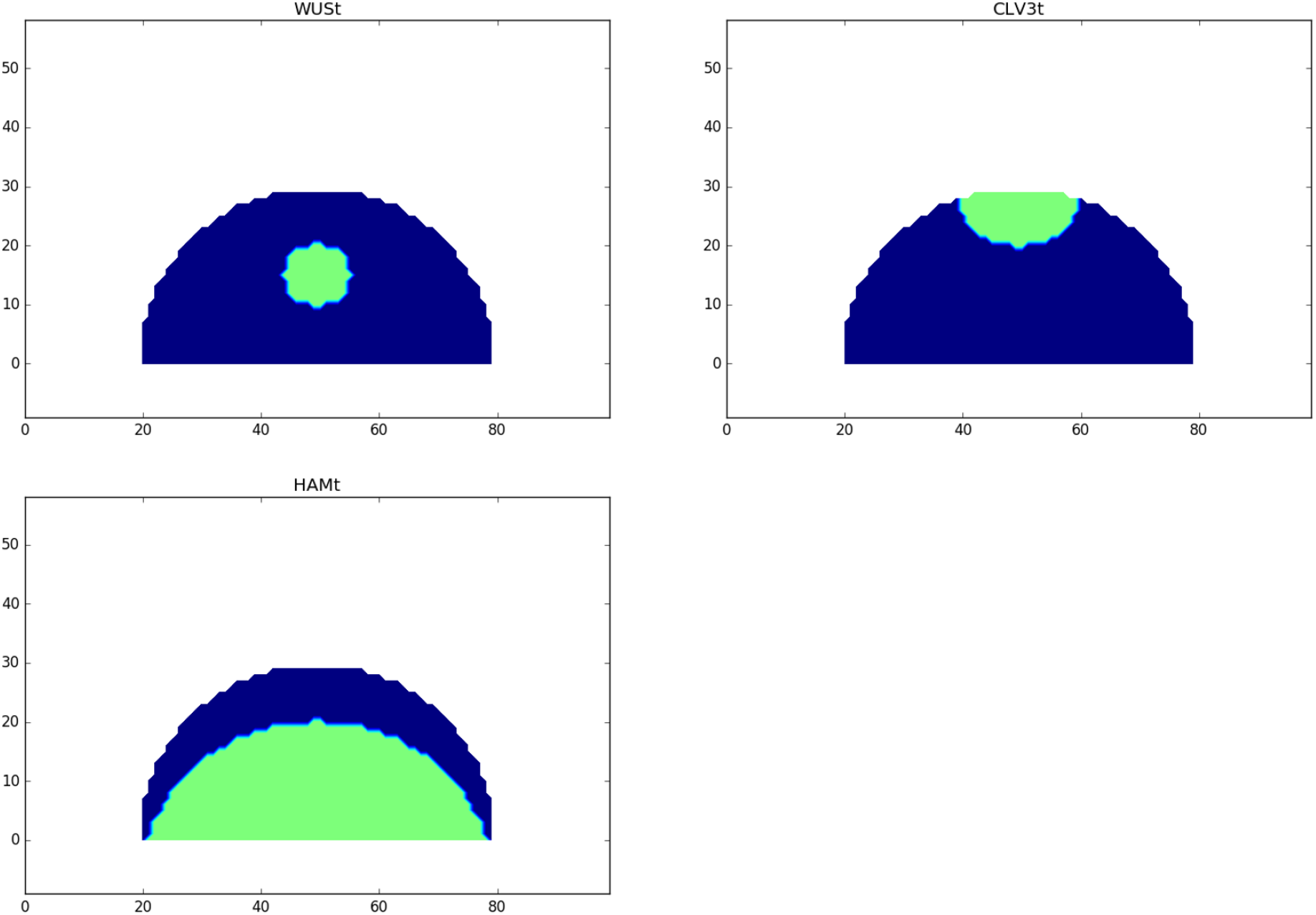
Optimisation target domains for WUS, *CLV3* and *HAM.*

